# Learning proteomic disease trajectories with flow matching

**DOI:** 10.64898/2026.07.08.737311

**Authors:** Erik Hartman, Christofer Karlsson, Johan Malmström

## Abstract

High-throughput proteomics has enabled detailed characterization of molecular states across health and disease. However, biological systems are inherently dynamic and methods for reconstructing continuous proteome changes remain limited. Here, we introduce proteome velocity, a framework for inferring continuous proteome trajectories from cross-sectional or sparsely sampled proteomics data using flow matching, in which a neural network learns velocity fields over proteome space. Proteome velocity estimates how rapidly and in which direction protein abundances change along a biological progression, such as disease. In mouse sepsis, covariate-conditioned velocity models resolved tissue- and pathogen-specific proteome trajectories and identified inflammatory proteins with distinct temporal activation patterns across infection routes and organ systems. In clinical COVID-19 plasma proteomes, inferred trajectories separated into distinct velocity programs associated with disease severity. These results show how generative trajectory models can transform cross-sectional proteomics data into interpretable, protein-resolved representations of molecular progression.

## 1 Introduction

Proteomics has emerged as a foundational technology for understanding human biology and disease. Recent advances in mass spectrometry have enabled the quantification of thousands of proteins across large cohorts, tissues, and increasingly individual cells, generating opportunities to characterize molecular phenotypes at scale. These developments have accelerated efforts to define molecular signatures of disease, identify biomarkers, and establish proteome-based taxonomies of biological systems [1].

Central to many of these efforts is the concept of the proteome state [2]. Across biomedical research, healthy and diseased tissues or distinct cellular phenotypes are frequently represented as discrete states. This conceptual framework has been successful, providing the basis for disease classification, patient stratification, and the identification of molecular correlates of biological function.

Despite this utility, a state-based view of the proteome has important limitations. Many biological processes are best understood as trajectories rather than discrete states. Cellular differentiation, tissue maturation, immune activation, infection, therapeutic response, aging, injury, and recovery all involve coordinated molecular changes that unfold over time. However, proteomic experiments typically measure samples at isolated time points or disease stages. Consequently, most proteomics analyses partition samples into groups and identify proteins that differ in abundance or summarize each protein by a single average slope over the progression axis. Although differential abundance analysis has proven effective for identifying disease-associated proteins and pathways [3], it imposes a simplified view on systems that are inherently dynamic and non-linear. When a biological progression is reduced to a series of endpoint comparisons, information about the direction, rate, and structure of molecular change is largely lost.

This challenge has motivated the development of computational trajectory inference methods in single-cell biology [4–10]. A wide range of approaches, including graph- and manifold-based representations, ordinary differential equations (ODEs), and optimal transport (OT) frameworks, have been developed to reconstruct developmental and cellular dynamics from cross-sectional transcriptomic measurements [4, 11–14]. Despite their methodological diversity, these approaches share a common objective of recovering continuous biological processes from sparsely observed molecular snapshots. More recently, advances in machine learning and generative modeling have introduced new frameworks for learning trajectories by modelling dynamical systems with neural networks [15–22].

Among these, flow matching has emerged as a powerful and computationally efficient framework for learning continuous transformations between probability distributions [15, 16, 18]. The goal of flow matching is to learn a time-dependent velocity field whose associated flow transports samples from a source distribution to a target distribution. Originally developed for generative modeling, the framework provides a natural representation of biological progression, where populations of molecular states evolve continuously over time. Recent studies have demonstrated the utility of flow matching for trajectory inference in single-cell transcriptomics, enabling the reconstruction of developmental and cellular transitions from population-scale measurements [17, 19, 23–25]. However, the application of trajectory inference to proteomics and beyond the single-cell scale remains largely unexplored.

Here, we utilize flow matching as a framework for proteomic trajectory inference at the organism level. Central to our approach is the concept of proteome velocity, defined as the local rate and direction of protein abundance change along an inferred trajectory. While differential abundance analysis quantifies the net difference between two molecular states, proteome velocity describes how those differences accumulate throughout a biological process. We first establish the theoretical framework for learning proteome trajectories using flow matching and then apply it to two distinct disease settings including tissue- and infection-model-specific sepsis progression in mice, and clinical COVID-19 severity progression in patients. Together, these analyses demonstrate how generative trajectory models can reveal dynamic features of proteome organization that are not accessible through conventional state-based analyses.

## 2 Flow matching for proteomic trajectories

To infer continuous proteomic dynamics from cross-sectional measurements, we adapted flow matching as a framework for proteomic trajectory inference. Rather than treating samples as isolated observations, flow matching models biological progression as a continuous transformation between distributions of molecular states. The method learns a time-dependent velocity field that describes how populations of samples move through a high-dimensional molecular space over the course of a biological process (**Fig. 1a-b**).

**Figure 1:**
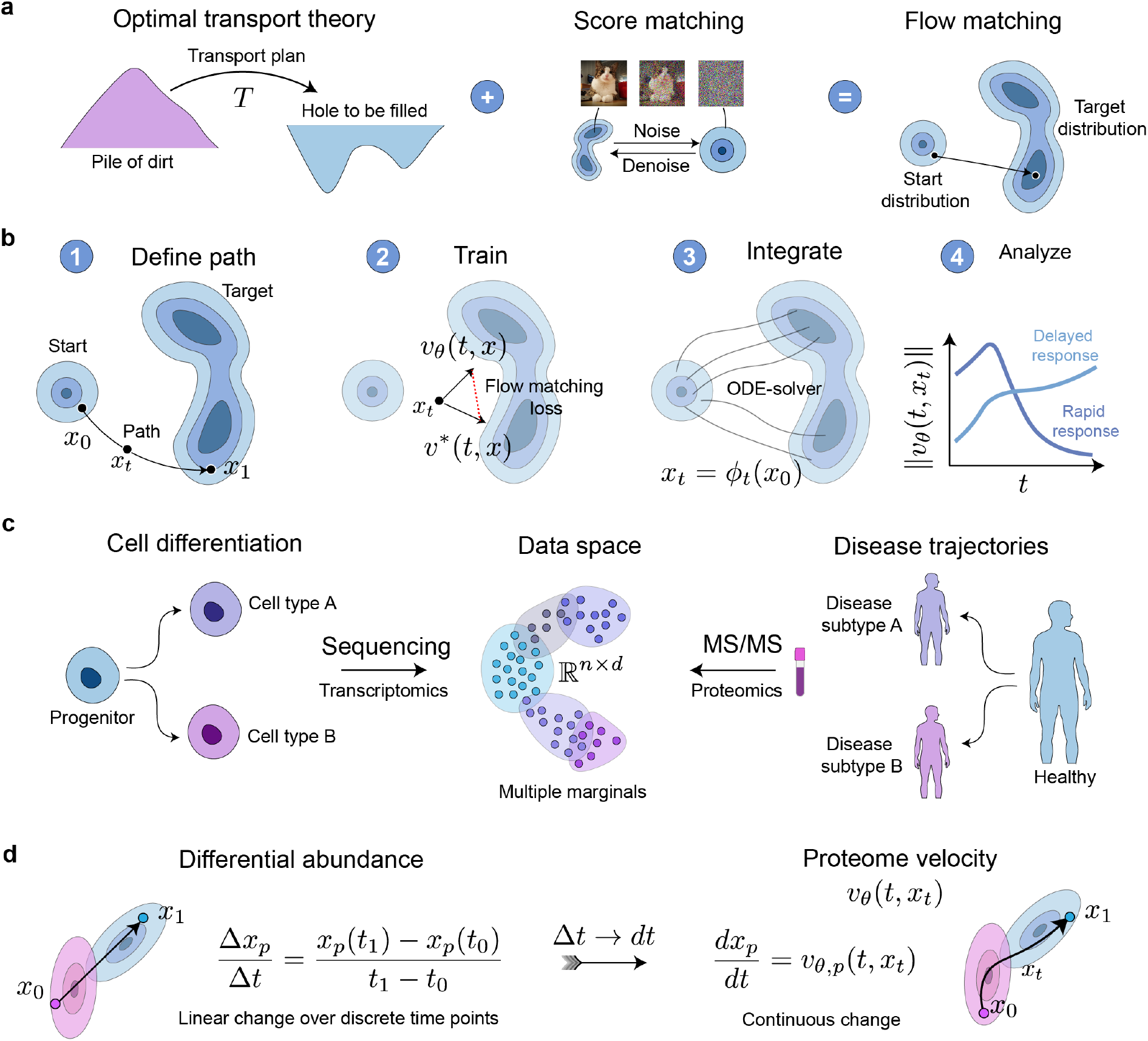
Flow matching for proteomic trajectory inference. **a** Flow matching links ideas from optimal transport and score-based generative modeling, while training can be performed without simulation. Optimal transport defines couplings between distributions, while continuous generative models describe how samples move through state space. Flow matching learns a velocity field whose flow transports one empirical distribution into another, combining the two paradigms. **b** The general workflow for using flow matching for trajectory inference begins with defining a path between source and target proteomic states, followed by training a model *v*_*θ*_(*t, x*) to predict the target velocity *v*^*∗*^(*t, x*) along that path. The model can then be used to integrate the learned flow *x*_*t*_ = *ϕ*_*t*_(*x*_0_), generating trajectories that can be analyzed to identify dynamic proteins and trajectory patterns. **c** Trajectory inference is well established in single-cell transcriptomics, where cells are ordered along developmental paths, represented as multiple marginals of an underlying developmental distribution. Proteomics measurements also place samples in a high-dimensional molecular space, but ordered processes such as disease progression are often analyzed as discrete group comparisons rather than continuous trajectories. **d d** There is a direct connection between the learned velocity field and conventional summaries of protein change. Differential abundance corresponds to a finite difference between states, and linear regression estimates an average slope over the progression axis, whereas proteome velocity is the corresponding local derivative, retaining information about the timing, magnitude and direction of protein-wise change along the modeled trajectory.

In proteomics, each sample can be represented as a point in a high-dimensional space defined by protein abundances. When samples are ordered along a temporal or pseudotemporal axis, flow matching learns how proteomes evolve through this space and reconstructs trajectories linking early and late molecular states (**Fig. 1c**). The learned velocity field provides a local estimate of the rate and direction of abundance change for every protein at every point along the trajectory. We refer to this protein-wise velocity field as proteome velocity (**Fig. 1d**).

Whereas differential abundance and linear regression quantify the constant difference between molecular states, proteome velocity describes how those differences accumulate throughout a biological process (see Supplementary information). This framework therefore enables continuous characterization of molecular progression and provides protein-resolved estimates of change that are not accessible through endpoint comparisons alone.

## 3 Inferring tissue- and model-specific disease trajectories in sepsis

Sepsis is a life-threatening syndrome caused by infection and characterized by a dysregulated systemic inflammatory response that can progress to organ failure and death [26]. Previous work in murine sepsis models has shown that disease progression is accompanied by a pronounced and time-dependent increase in acute-phase and inflammatory response proteins across multiple organs and in plasma. This inflammatory response is paralleled by the appearance of tissue-specific proteins in circulation, suggesting that sepsis induces widespread organ injury characterized by inflammatory activation within tissues and leakage of intracellular proteins into plasma [27]. Given the dynamic nature of these processes, we reasoned that proteome velocity could provide a framework for resolving how inflammatory proteome remodeling unfolds across tissues and infection routes during disease progression.

As a starting point, we analyzed a previously published sepsis resource comprising 1,578 proteome maps from human samples and seven animal models of sepsis. For the present analysis, we focused on 704 time-resolved proteome maps from murine models spanning seven tissues and cell types including heart, kidney, leukocytes, liver, lung, plasma, and spleen, covering 10,181 identified proteins in total (**Fig. 2a**). The dataset included six pathogen-route combinations, intranasal *Klebsiella pneumoniae* (IN-Kpn), intranasal *Streptococcus pneumoniae* (IN-Spn), intraperitoneal *Escherichia coli* (IP-Eco), intraperitoneal *Streptococcus pyogenes* (IP-Spy), intravenous *Staphylococcus aureus* (IV-Sau) and subcutaneous *Streptococcus pyogenes* (SC-Spy). Samples were collected at multiple time points throughout disease progression and aligned to a normalized disease-time axis to account for differences in progression rates across sepsis models.

**Figure 2:**
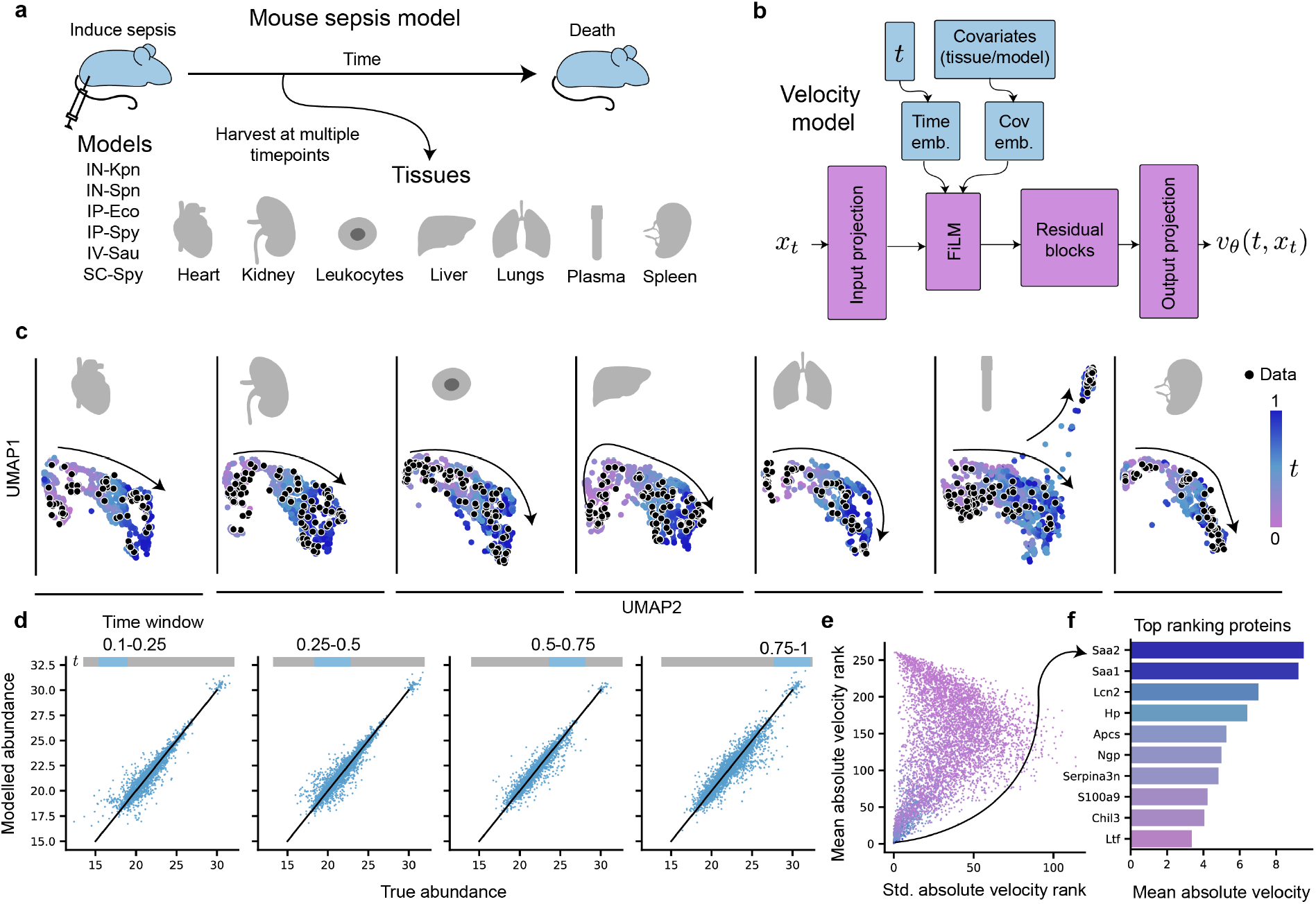
Flow matching model for mouse sepsis proteomics. **a** Experimental design for the mouse sepsis dataset. Six sepsis models representing different combinations of pathogen and infection route were sampled across multiple disease time points and seven tissues or cell types. Disease time was normalized to account for different model-specific progression kinetics. **b** Overview of the proteome velocity model architecture. The model receives proteome state *x*_*t*_, normalized disease time *t* and tissue/model co-variates, embeds the covariates through FiLM conditioning, and outputs the predicted proteome velocity *v*_*θ*_(*t, x*_*t*_). **c** UMAP projections of generated validation trajectories for each tissue. Trajectory points are colored according to normalized disease time and observed samples are shown as black points. **d** Comparison of modeled and observed protein abundances (True abundance) across four disease-time intervals. **e** Stability of protein velocity rankings across five independently trained models. Proteins with high mean absolute velocity ranks and low rank variability represent robust dynamic features of disease progression. **f** Top proteins based on mean absolute velocity across generated validation trajectories.

To focus on proteome remodeling shared across tissues and sepsis models, we restricted the analysis to proteins quantified across all tissue-model combinations, yielding 262 proteins measured in all 704 samples. Functional enrichment analysis showed that these proteins were predominantly associated with platelet and neutrophil degranulation, complement and coagulation cascades, and acute inflammatory responses (**Supplementary Fig. S1**). Protein abundances were mean-centered within each tissue–model combination to reduce systematic offsets while preserving the shared protein scale before training a covariate-conditioned flow-matching model. The model used the 30 largest principal components, normalized disease time, tissue identity, and sepsis model as inputs and learned a proteome velocity field describing local abundance changes throughout disease progression. The model was implemented as a residual network [28] with feature-wise linear modulation (FiLM [29]) conditioning, and its output was the predicted proteome velocity vector (**Fig. 2b**). Conditioning on covariates allowed the velocity model to leverage information across tissues and models while preserving context-specific structure.

To ensure that trajectories reflected biological progression, source-target pairs were sampled only in the forward time direction, primarily between adjacent disease stages. Five independent models were trained using different random initializations. Integrating the learned velocity fields from validation samples accurately reconstructed the observed distributions of disease states across tissues and sepsis models (**Fig. 2c**). Predicted and observed protein abundances remained highly concordant throughout progression (mean Pearson *r ≈* 0.95; **Fig. 2d**), indicating that the learned velocity field captures the dominant proteomic changes associated with disease progression.

The highest-ranking proteins were highly consistent across model-replicates (**Fig. 2e-f**). The proteins with the largest mean absolute velocities included serum amyloid A-2 (Saa2), serum amyloid A-1 (Saa1), neutrophil gelatinase-associated lipocalin (Lcn2), haptoglobin (Hp), serum amyloid P-component (Apcs), neutrophilic granule protein (Ngp), serine protease inhibitor A3N (Serpina3n), protein S100-A9 (S100a9), chitinase-like protein 3 (Chil3) and lactotransferrin (Ltf). Four of these proteins, Saa1, Saa2, Hp, and Apcs, are classical acute-phase proteins produced primarily by the liver in response to inflammatory cytokines. Others, including Lcn2, S100a9, Ngp, Chil3, and Ltf, are associated with neutrophil activation, antimicrobial defense, and innate immune responses. Serpina3n is similarly induced during inflammation and tissue injury and functions as a protease inhibitor that limits inflammatory damage.

The high velocities of these proteins indicate that their abundances change rapidly along the inferred disease trajectories, consistent with their established roles as early responders to infection and inflammation. Unlike constitutively expressed proteins that remain relatively stable, acute-phase and innate immune proteins can be induced within hours of infection, resulting in large and coordinated shifts in abundance. Their strong velocity signals across tissues and plasma therefore reflect the extensive proteome remodeling that accompanies progression toward severe inflammatory states.

## 4 Proteome velocity resolves tissue- and pathogen-specific inflammatory dynamics

We next used the learned velocity field to determine where and when proteomic remodeling was most pronounced during sepsis progression. Integrating trajectories and stratifying by tissue revealed substantial differences in overall proteome velocity in different sample types. Plasma exhibited the highest mean absolute velocity, followed by leukocytes, whereas the lungs had the lowest overall velocity (**Fig. 3a**). Stratification by sepsis model further revealed that most sepsis trajectories were characterized by a declining proteome velocity over time, showing that the largest proteomic changes among the selected proteins occurred early after infection. Notable exceptions were the subcutaneous and intranasal models (SC-Spy and IN-Kpn), which showed more delayed or sustained velocity profiles.

**Figure 3:**
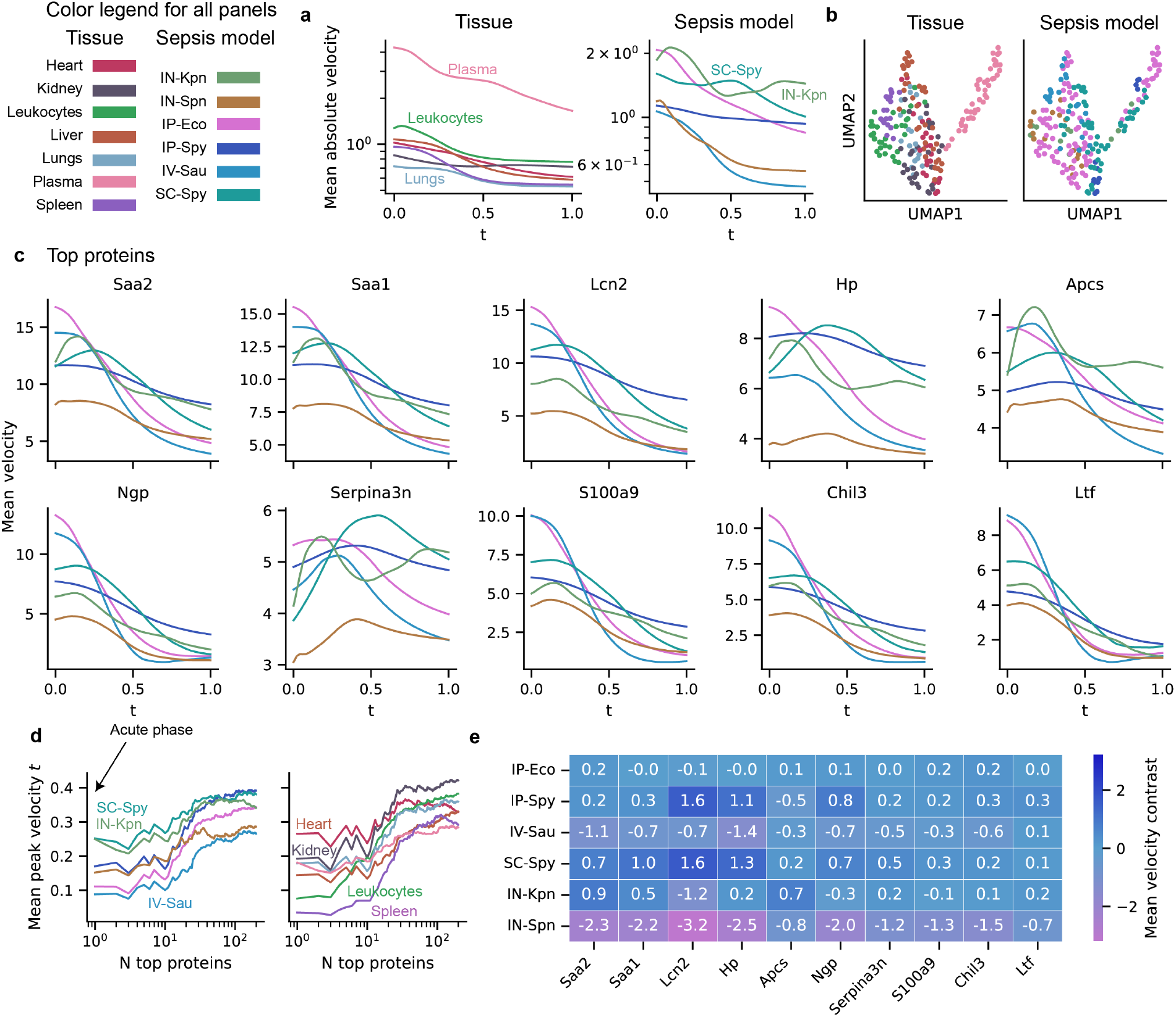
Tissue- and model-specific proteome velocity during mouse sepsis progression. **a** Mean absolute proteome velocity over normalized disease time, stratified by tissue (left) and by sepsis model (right). **b** UMAP projection of protein-wise velocity profiles. Each dot represents a sample and is colored by tissue or sepsis model. **c** Mean velocity profiles of the ten most dynamic proteins colored by sepsis model. **d** To assess whether protein velocity peak timing was associated with velocity magnitude, proteins were ranked by absolute mean velocity and progressively included using increasing rank cutoffs. For each cutoff, the timing of the mean protein velocity peak was averaged. The plots show the number of included proteins across peak times, stratified by sepsis model (left) and tissue (right). **e** To identify infection-model-specific deviations from the average velocity profile, protein-wise contrast profiles were computed by subtracting the mean velocity profile from each individual profile. The heatmap shows the mean velocity contrast for the top proteins, stratified by sepsis model. For all aggregated velocity profiles, confidence bands are not shown for visual clarity.

Projection of protein-wise velocity trajectories using UMAP demonstrated clustering by both tissue and sepsis model, suggesting that velocity captures sepsis model-specific disease programs rather than simply reflecting the magnitude of systemic inflammation (**Fig. 3b**). Inspection of individual proteins revealed marked differences in the timing of peak velocity across sepsis models. For most proteins, velocities peaked early, especially in the IP and intravenous (IV) models, whereas subcutaneous and intranasal models displayed delayed responses (**Fig. 3c**). The delayed response observed in SC-Spy is consistent with the additional time required for a localized infection to disseminate and trigger systemic sepsis. Across the highest velocity proteins, the estimated timing of peak velocity followed an approximate IV *→* IP *→* IN *→* SC progression, matching the expected kinetics of pathogen dissemination from direct bloodstream exposure to progressively slower local infection routes.

Among the top velocity proteins, Serpina3n, the mouse orthologue of alpha-1-antichymotrypsin, showed a distinct temporal profile, with a delayed velocity peak across all sepsis models (**Fig. 3c**). Serpina3n is a protease inhibitor, and its delayed induction relative to classical acute-phase proteins is consistent with a secondary protective or regulatory response that follows the initial inflammatory burst [30].

When progressively lower-ranked proteins were included, the average timing of peak velocity shifted later in disease progression. This indicates that the highest-velocity proteins tend to respond earliest, whereas proteins with lower velocity magnitudes show more delayed dynamics (**Fig. 3d**). A similar pattern emerged when trajectories were stratified by tissue (**Fig. 3d**). The spleen and leukocytes, which play central roles in initiating and coordinating the inflammatory response, exhibited the earliest and most rapid proteomic changes. In contrast, the heart and kidney showed substantially delayed responses, consistent with these organs acting primarily as targets of systemic inflammation rather than major sites of immune activation. Together, these findings demonstrate that proteome velocity captures both pathogen-specific differences in disease progression and biologically meaningful variation in the timing of tissue responses.

To identify pathogen- and route-specific responses, we computed velocity contrasts by calculating the global mean velocity profile for all models followed by subtracting the global mean velocity profile from each model-specific profile. In this analysis, a positive contrast indicates that a protein’s velocity was higher than the average profile at the corresponding disease time, revealing clear infection-specific deviations among the highest-velocity proteins (**Fig. 3e**). For example, IN-Spn showed consistently lower velocity contrasts, whereas both the intraperitoneal and subcutaneous *S. pyogenes* sepsis models showed elevated velocity contrasts for Lcn2 and Hp. Lcn2 has previously been reported to be induced in a *S. pyogenes*-associated inflammatory setting [31], and Hp has been implicated as a potential protective factor in streptococcal disease [32]. These findings demonstrate that proteome velocity retains the protein-level interpretability of conventional differential abundance analyses while additionally providing temporal ordering and model-specific information about the dynamics of disease progression.

## 5 Proteome velocity extends conventional summaries of protein change

We next compared proteome velocity with conventional summaries of protein change. We first performed a differential abundance analysis by comparing samples collected before and after the midpoint of normalized disease time (**Fig. 4a**). This endpoint contrast was strongly correlated with mean protein velocity estimated by flow matching (Spearman *ρ* = 0.97, *p <* 0.001), supporting the interpretation of proteome velocity as a local, time-resolved analogue of differential abundance.

**Figure 4:**
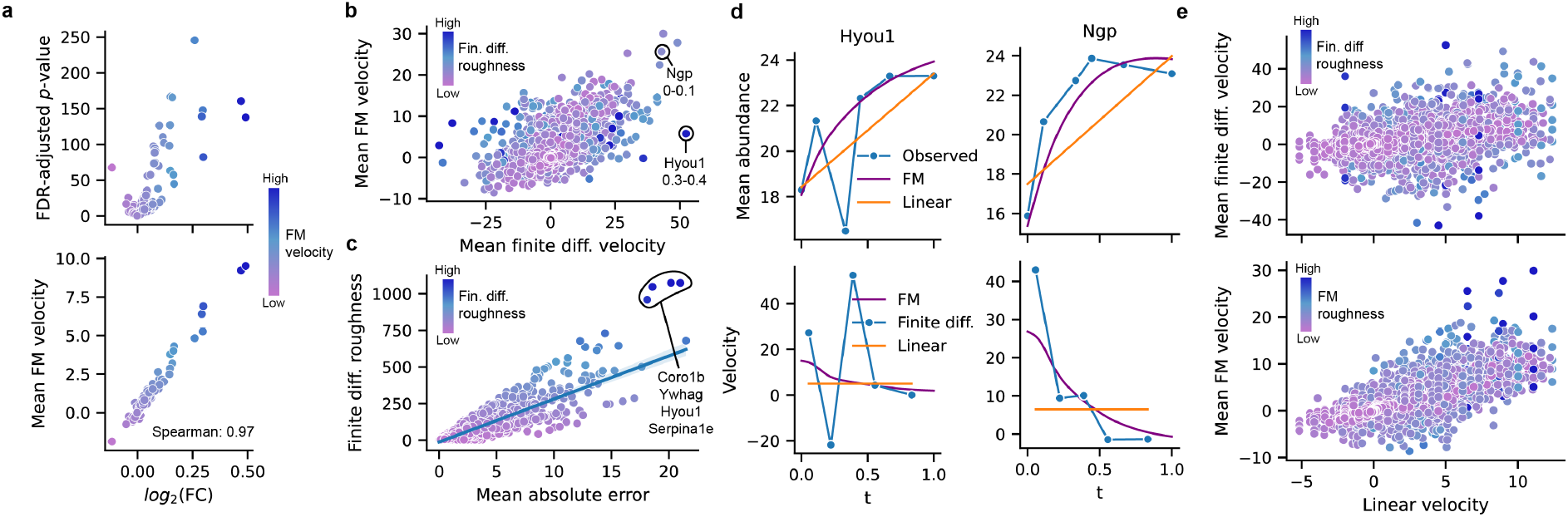
Relationship between differential abundance, finite-difference velocity, linear velocity and flow-matching velocity. **a** Differential abundance analysis comparing samples before and after the normalized-time midpoint (upper). Mean flow-matching velocity was strongly correlated with the resulting *log*_2_ fold change (Spearman *ρ* = 0.97, lower). **b** Comparison of empirical finite-difference velocity and flow-matching velocity across protein, tissue, model and time-interval combinations. Points are colored by roughness of the finite-difference velocity profile. **c** Relationship between finite-difference roughness and mean absolute error between empirical finite-difference and flow-matching velocities. Each point represents one protein–tissue–model trajectory. Points are colored by finite-difference roughness, and the line shows the linear regression estimate. **d** Example abundance and velocity profiles for Hyou1 (left) and Ngp (right) in IP-Eco plasma. Hyou1 shows a sharp finite-difference fluctuation that is smoothed by flow matching, whereas Ngp shows more concordant finite-difference and flow-matching dynamics. **e** Comparison of the linear-model velocity with empirical finite-difference velocity (upper) and flow-matching velocity (lower).

We next compared flow-matching velocities with empirical finite-difference estimates of protein dynamics. For each protein, tissue, and sepsis model, we first summarized the observed trajectory by calculating the mean log-abundance at each sampled time point. Empirical velocities were then computed between adjacent time points as the change in mean log-abundance divided by the corresponding change in normalized disease time. Thus, each empirical velocity represents the slope of the straight line connecting two consecutive observed time points. These finite-difference estimates provide a direct empirical reference for the velocity learned by the flow-matching model, since the model is trained to represent changes in protein abundance over disease time. To make the two quantities comparable, flow-matching velocities were averaged over the same time intervals used for the finite-difference calculation. Overall, the empirical and flow-matching velocities showed agreement (*R*^2^ = 0.33), indicating that the model captured a substantial component of the observed temporal dynamics. However, several proteins showed large empirical finite-difference velocities but much smaller flow-matching velocities (**Fig. 4b**).

We reasoned that these discrepancies could arise when the empirical trajectory contains abrupt local fluctuations. Such fluctuations can produce large finite-difference velocities over individual intervals, even if the broader temporal trend is relatively smooth. To quantify this behavior, we defined velocity roughness as the summed absolute change in empirical velocity between adjacent intervals. This measure captures how strongly the finite-difference velocity profile changes from one interval to the next. Protein trajectories with higher roughness showed larger mean absolute disagreement between finite-difference and flow-matching velocities (Pearson *r* = 0.88, **Fig. 4c**), suggesting that irregular empirical dynamics were smoothed by the flow-matching model rather than captured exactly.

For the proteins with the largest roughness, including Hypoxia up-regulated protein 1 (Hyou1), 14-3-3 protein gamma (Ywhag), Coronin-1B (Coro1b) and Alpha-1-antitrypsin 1-5 (Serpina1e), the finite-difference estimates were driven by sharp local changes between neighboring time points, whereas the flow-matching model emphasized smoother temporal trends across disease progression, as exemplified by Hyou1 (**Fig. 4d**). Upon further inspection of the abundance profiles for these proteins, it was found that the jagged abundance profile for these outlying proteins was due to a combination of batch differences and uneven sampling frequency (**Supplementary Fig. S2**). These examples suggest that flow matching captures robust temporal trends while smoothing over abrupt finite-difference fluctuations that may arise from sparse sampling, measurement noise or biological variability.

To investigate the extent to which the flow matching model captured non-linear dynamics, we compared it with a linear model fitted separately for each protein, tissue and sepsis model, using normalized disease time as the sole predictor of protein abundance. The resulting slope represents a single constant velocity over the entire disease trajectory. As expected, this linear velocity provided the most heavily smoothed description of temporal change and correlated poorly with the observed finite-difference velocity (*R*^2^ = *−*3.1, **Fig. 4e**). The linear model generally underestimated velocity magnitudes and failed to capture context-dependent changes in trajectory shape. The largest discrepancies between linear and flow-matching velocities occurred for proteins exhibiting highly dynamic velocity profiles, highlighting the inability of a constant-velocity approximation to represent nonlinear disease progression. As such, proteome velocity provides an interpretable and biologically meaningful representation of protein dynamics that extends conventional analysis methods into a continuous and dynamic framework.

## 6 Inferring severity trajectories in patients with COVID-19

An ordered progression axis does not necessarily have to represent physical time. Instead, any biologically meaningful ordered variable can serve as a trajectory coordinate, provided that the resulting trajectories are interpreted as progression paths rather than physical time courses. To evaluate whether proteome velocity could capture disease progression in such a setting, we applied flow matching to plasma proteomes from patients with COVID-19.

In the original study, patients were categorized according to the WHO Covid-19 severity scale and profiled using mass spectrometry-based plasma proteomics [33]. The samples span WHO grades 3-7 and this severity gradient was used as an ordered pseudotemporal axis. After filtering for proteins quantified in all samples, 94 proteins remained for downstream analysis. Protein abundances were log_2_ transformed and centered within each cohort to reduce dataset-specific offsets, and mapped onto a normalized severity axis based on WHO grade. We then trained a flow-matching model to infer trajectories connecting lower- and higher-severity disease states (**Fig. 5a**). The model architecture and training procedure were otherwise identical to those used in the sepsis analysis, except that no tissue or infection-model covariates were included.

**Figure 5:**
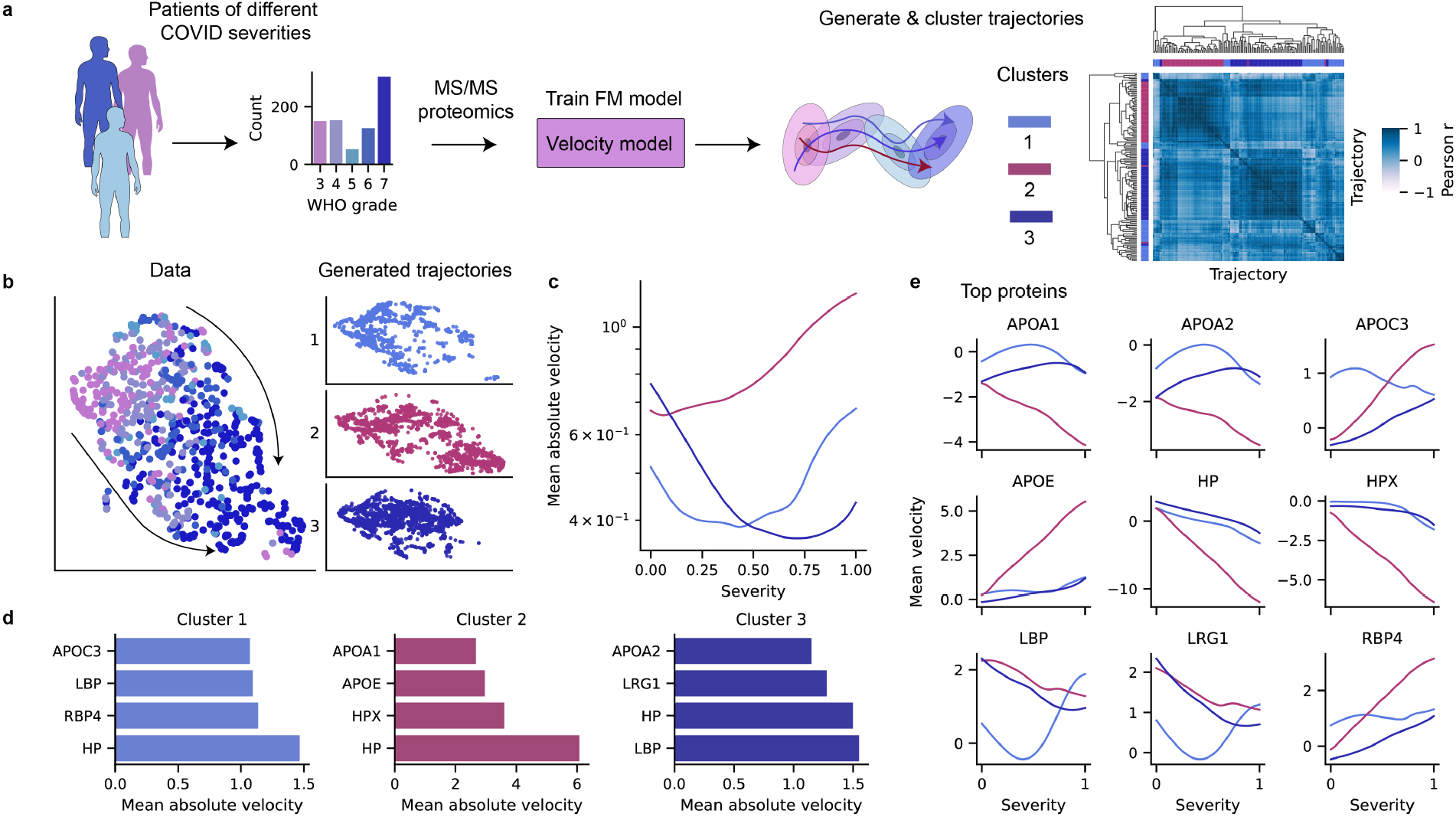
Proteome velocity along COVID-19 severity trajectories. **a** Workflow for COVID-19 severity modeling. Plasma proteomes from patients across WHO severity grades 3–7 were used to train a flow matching model on a normalized severity axis. Generated validation trajectories were clustered by velocity-profile similarity, yielding three trajectory clusters. **b** UMAP projection of observed data and generated trajectories. Generated trajectories separated into three clusters with distinct locations in proteome space. **c** Mean absolute proteome velocity over severity for each trajectory cluster. Cluster 1 showed a U-shaped velocity profile over severity, whereas cluster 3 showed a declining velocity, and cluster 2 showed increasing velocity. **d** Top proteins by mean absolute velocity within each trajectory cluster. **e** Cluster-stratified velocity profiles for top dynamic proteins over the normalized severity axis, showing that severity-associated protein changes differed in magnitude, direction and timing between trajectory clusters. For all aggregated velocity profiles, confidence bands are not shown for visual clarity.

Although no covariates were included in the model training, integrating the learned velocity field from validation samples revealed heterogeneity in the disease trajectories. To compare the inferred proteomic velocity patterns across samples, we computed pairwise correlations between each sample’s velocity profile across proteins. Agglomerative clustering of this sample-by-sample correlation matrix separated the cohort into three major groups, which we refer to as three velocity programs (**Fig. 5a**). These three clusters occupied distinct regions of trajectory space when projected using UMAP (**Fig. 5b**) and exhibited different velocity profiles (**Fig. 5c**). These distinct velocity profiles can be thought of as canyons in the learned proteome velocity field, representing distinct paths transporting samples from low to high severity. In contrast to the predominantly declining velocity profiles observed during sepsis progression, cluster 1 had a U-shaped mean absolute velocity pattern, cluster 2 showed progressively increasing velocity with severity, and cluster 3 had a declining profile (**Fig. 5c**). These different patterns suggest that increasing clinical severity is not simply associated with a linear attenuation or amplification of the same proteomic program but are associated with distinct proteomic states.

The top proteins by mean absolute velocity differed between the three clusters (**Fig. 5d**). Cluster 2 was dominated by haptoglobin (HP), hemopexin (HPX), apolipoprotein E (APOE), apolipoprotein A-I (APOA1) and apolipoprotein A-II (APOA2), with HP showing the largest velocity. This observation is consistent with the original study, which reported reduced HP and HPX levels in severely ill patients, particularly those undergoing renal replacement therapy (RRT) or extracorporeal membrane oxygenation (ECMO). Moreover, the apolipoproteins showed heterogeneous dynamics across clusters, recapitulating findings from the source study showing that apolipoprotein patterns varied with covariates like age [33].

Cluster 1 was likewise characterized by high HP velocity but additionally showed strong contributions from retinol-binding protein 4 (RBP4), a protein previously associated with COVID-19 susceptibility, hospitalization, and disease severity [34]. In contrast, lipopolysaccharide-binding protein (LBP) was the dominant protein in cluster 3 and also ranked highly in cluster 1, despite not being highlighted in the original study. Although LBP is classically associated with responses to bacterial lipopolysaccharide (LPS), elevated LBP levels have been linked to post-COVID-19 syndrome [35], and interactions between the SARS-CoV-2 spike protein and LPS have been reported [36]. Examination of cluster-specific velocity profiles further revealed substantial differences in both the magnitude and direction of protein changes across disease severity (**Fig. 5e**).

Together, these results demonstrate that proteome velocity can recover heterogeneous progression programs from ordered clinical datasets, even when the trajectory axis reflects disease severity rather than physical time. Combined with the sepsis analyses, these findings establish flow matching as a general framework for extracting dynamic, protein-resolved representations of biological progression from cross-sectional and sparsely sampled proteomic data.

## 7 Discussion

Proteomics has traditionally been interpreted through the notion of molecular states, where biological systems are represented as discrete points within a high-dimensional proteomic landscape. Here, we show that flow matching provides a natural framework for extending this view toward non-linear continuous proteome dynamics. By learning velocity fields over proteome space, our approach enables reconstruction of molecular trajectories from cross-sectional or sparsely sampled proteomics data and provides protein-resolved estimates of how proteomes change along a biological progression axis. Whereas differential abundance analysis quantifies the net difference between molecular states, proteome velocity describes how those differences accumulate throughout a biological process. The correspondence between differential abundance and inferred velocity demonstrates that the framework captures biological signals while additionally providing information about the timing and direction of molecular change.

The sepsis analysis illustrates the value of this perspective. Although acute-phase and inflammatory proteins dominated the response across sepsis models, their velocity profiles differed substantially. Intravenous and intraperitoneal infections induced rapid early proteome remodeling, whereas the peak velocity of many highly dynamic proteins was delayed in the subcutaneous and intranasal models. This pattern likely reflects differences in pathogen dissemination kinetics, with localized infections requiring additional time before eliciting a systemic inflammatory response. Definitive validation of this hypothesis would require denser temporal sampling during the transition from local to systemic infection, which remains challenging in vivo.

The proteins exhibiting the highest velocities were primarily associated with the acute-phase response, neutrophil activation, antimicrobial defense, and innate immunity. Classical acute-phase proteins, including Saa1, Saa2, Hp, and Apcs, are predominantly synthesized by the liver and rapidly released into circulation following inflammatory stimulation. Their high velocities therefore reflect the extensive hepatic reprogramming that accompanies the acute-phase response. In contrast, the delayed velocity peak observed for the protease inhibitor Serpina3n distinguished it from the classical acute-phase proteins and is consistent with a later protective response aimed at limiting inflammation-induced tissue injury [30]. These temporal relationships are difficult to resolve using conventional endpoint comparisons or constant slopes but emerge when disease progression is represented as a continuous trajectory.

The observation that plasma, leukocytes, and spleen exhibited the highest overall proteome velocities is likewise consistent with their central roles in coordinating the systemic response to infection. Plasma serves as the principal transport medium for circulating acute-phase proteins and therefore integrates inflammatory signals originating from multiple organs. Circulating leukocytes undergo extensive transcriptional and proteomic reprogramming in response to pathogen exposure, while the spleen acts as a major reservoir and activation site for innate immune cells. Consequently, many of the proteins displaying the highest velocities, including Lcn2, S100a9, Ngp, Chil3, and Ltf, are linked to neutrophil activation and innate immune responses and would be expected to change most rapidly within these compartments. In contrast, organs such as the heart and kidney primarily experience the downstream consequences of systemic inflammation and therefore exhibit slower and less pronounced proteome remodeling. These findings suggest that proteome velocity captures not only the magnitude of molecular change but also the hierarchical organization of the host response, with immune and circulatory compartments acting as primary sites of rapid proteomic adaptation during sepsis.

The COVID-19 analysis shows that the same framework is not restricted to time-series experiments. By treating disease severity as an ordered pseudotemporal axis, we identified multiple velocity programs associated with progression toward severe disease. Importantly, these trajectories should not be interpreted as causal models of patient deterioration or as reconstructions of individual patient histories. Rather, they represent inferred paths through severity-associated proteomic states. Within this interpretation, the results suggest that increasing severity is not characterized by a single homogeneous proteomic program, but instead can arise through distinct velocity programs with different dominant proteins and velocity profiles.

Several limitations should be considered. First, current proteomic datasets remain substantially smaller than single-cell transcriptomic datasets. Consequently, trajectory models applied to proteomics must operate in a regime where sample size and sampling frequency is the primary constraint. In this study, we addressed this challenge by using a simple velocity model, early stopping, repeated training with different random seeds and downstream analyses focused on high-velocity proteins that were robust across model replicates. Nevertheless, future work should systematically evaluate alternative flow-matching formulations, optimal-transport and Schrödinger-bridge couplings, neural ODE architectures, and uncertainty-aware trajectory models specifically tailored to proteomic data. Second, the sepsis analysis was restricted to the 262 proteins quantified across all tissues, sepsis models, and time points. While this reduced the dimensionality of the data, ensured a complete dataset and enabled direct comparison across biological contexts, it biased the analysis toward highly abundant and consistently detected proteins, particularly acute-phase and inflammatory mediators. Consequently, the inferred velocity field primarily captures systemic proteome remodeling, whereas tissue-specific, low-abundance, or transiently expressed proteins are underrepresented. Future methods that explicitly accommodate missing values may enable trajectory inference across a broader and more tissue-specific fraction of the proteome.

More broadly, our findings suggest that many proteomic studies may benefit from a trajectory-centered perspective. Biological processes such as disease progression, treatment response, recovery, development, aging, and cellular differentiation are inherently dynamic, yet are often analyzed as collections of discrete groups. Whenever a biologically meaningful ordering exists, trajectory models provide an opportunity to recover temporal structure that is otherwise obscured by state-based analyses. The framework presented here is therefore applicable not only to longitudinal studies, but also to cross-sectional datasets in which samples can be ordered according to disease severity, developmental stage, treatment response, or other progression variables.

Advances in mass spectrometry, increasing sample throughput, and the rapid growth of bulk and single-cell proteomics repositories will provide new possibilities for developing trajectory-based analytical frameworks. As larger and more densely sampled cohorts become available, generative models will increasingly enable proteomics to move beyond static descriptions of molecular states toward quantitative models of molecular progression. Such developments will catalyze the transition from proteome states to proteome dynamics and provide a more predictive understanding of biological processes across health and disease.

## 8 Methods

### 8.1 Sepsis data preprocessing

Mouse sepsis proteomics data were assembled from tissue-specific LC-MS/MS quantification tables and the corresponding sample metadata. The analysis included heart, kidney, leukocytes, liver, lungs, plasma and spleen samples from six sepsis models: IN-Kpn, IN-Spn, IP-Eco, IP-Spy, IV-Sau and SC-Spy. Protein groups with ambiguous identifiers were removed, and the tissue-specific matrices were then merged into a single sample-by-protein table. Proteins were retained only if they had complete measurements across the merged dataset used for the main analysis, resulting in 704 samples and 262 proteins.

The disease axis was the normalized hours post-infection variable supplied with the sample metadata. This variable maps model-specific disease progression to a shared interval from 0 to 1 and was used as the temporal coordinate throughout the sepsis analysis. Protein abundance values were treated as log-scale intensities. To reduce systematic tissue–model offsets while preserving the shared protein scale, each protein was centered within each tissue–model group and shifted back by the overall mean of the group means. The processed samples were split into training and validation sets using an 80/20 split stratified by binned disease time.

### 8.2 COVID-19 data preprocessing

COVID-19 plasma proteomics data were assembled from the Charité and Innsbruck cohorts described in the source study [33]. Ambiguous protein identifiers were removed, and proteins with missing values within either cohort were excluded before merging the two datasets. The Innsbruck severity annotation was renamed to the shared WHO grade field. The data contained WHO grade 0 values, which were treated as mislabels in light of the original study and recoded as grade 3. The final processed dataset contained 786 samples and 94 proteins, including 687 Charité samples and 99 Innsbruck samples.

Protein abundances were log_2_ transformed and corrected for dataset-specific offsets by subtracting the dataset-specific mean abundance for each protein and adding back the mean of the dataset means. WHO grades 3–7 were mapped to a normalized severity coordinate, so that grades 3, 4, 5, 6 and 7 corresponded to *t* = 0, 0.25, 0.5, 0.75 and 1, respectively. The COVID-19 model used this severity coordinate as a pseudotemporal axis. Samples were split into training and validation sets using an 80/20 split stratified by binned severity.

### 8.3 Pair sampling and flow-matching objective

For a source sample *x*_0_ at time *t*_0_ and a target sample *x*_1_ at a later time *t*_1_, an intermediate time *t* was sampled uniformly from the interval [*t*_0_, *t*_1_]. The corresponding intermediate state was defined by linear interpolation,

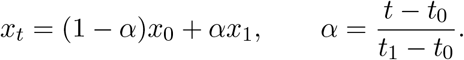

The target velocity for the pair was

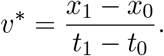

The flow matching model was trained to predict this velocity from the interpolated state and time by minimizing the mean squared error between *v*_*θ*_(*t, x*_*t*_) and *v*^*∗*^. Three interpolation times were sampled for each selected pair during loss evaluation.

Pairs were sampled only in the forward direction along the temporal or pseudotemporal axis. Samples were first grouped into discrete time buckets. In the sepsis analysis, pairs were sampled within the same tissue–model combination, whereas in the COVID-19 analysis pairs were sampled across the severity axis without additional covariates. Adjacent time-bucket transitions were sampled most frequently, with one-skipped-bucket transitions included to provide slightly longer local moves. Specifically, step sizes of 1 and 2 were sampled with probabilities 0.85 and 0.15. Group and transition sampling were balanced so that large covariate groups or densely sampled intervals did not dominate the objective. Target samples were selected with a small bias toward nearby states in proteome space. In each training epoch, 3000 source–target pairs were sampled and pairs were redrawn between epochs. This sampling strategy was used to reduce the influence of group-size imbalance and to emphasize local temporal transitions in relatively small datasets. This setup mimics multi-marginal flow matching with linear interpolation (L-MMFM) [19].

### 8.4 Velocity model architecture and training

For both datasets, the standardized protein matrix was reduced to 30 principal components, fitted on the training set only, as is common in trajectory inference workflows [6, 12]. This retained 87% of the variance in the COVID-19 data, and 77% of the variance in the sepsis data. The velocity model was trained in this reduced state space, and generated abundances and velocities were mapped back to protein space for downstream analyses.

The velocity field was represented by a residual neural network [28] trained in the 30-dimensional PCA-reduced proteomic state space, implemented in PyTorch [37]. The network takes the current reduced proteomic state, scalar time coordinate and, where applicable, covariates as input, and outputs a velocity vector in the same reduced state space. Time was encoded with sinusoidal features [38]. In the sepsis velocity model, tissue and sepsis model were included as categorical covariates; these covariates were embedded and combined with the time embedding to form a conditioning vector that modulated hidden representations through feature-wise affine conditioning (FiLM [29]). No covariates were used in the COVID-19 model. The network used an input projection layer, three residual blocks, hidden width 512, dropout 0.1, a 64-dimensional time embedding and 64-dimensional categorical covariate embeddings where applicable. Layer normalization and SiLU activation functions were used in the input projection and residual blocks. An output projection layer was appended after the residual blocks. The COVID-19 model contained approximately 2.0 million trainable parameters, whereas the sepsis model contained approximately 2.5 million parameters due to the additional covariate embeddings and FiLM conditioning layers, making both models easily trainable on an Apple Silicon M2 Max laptop.

For each dataset, five independent models were trained with different random seeds. Models were optimized with Adam [39] using a learning rate of 10^*−*4^ and weight decay of 10^*−*4^. The learning rate was reduced by a factor of 0.5 when the validation loss plateaued, and training was stopped early if the validation loss failed to improve by at least 10^*−*4^ for 10 epochs. Training was allowed to run for at most 100 epochs, and the model state with the best validation loss was retained. Minor run-to-run variation in generated trajectories can occur due to non-deterministic operations in the computational stack; downstream analyses were therefore based on aggregate patterns across independently seeded model replicates, and the biological interpretations are therefore robust.

### 8.5 Trajectory generation and downstream analysis

After training, trajectories were generated from samples at the start of the disease or severity axis. For each validation start sample with *t* = 0, the learned velocity field was integrated from *t* = 0 to *t* = 1 using the explicit midpoint second-order Runge-Kutta method with 50 time steps, yielding 51 trajectory states per generated path. The same procedure was also applied to training start samples for quality control, but all main downstream analyses used validation trajectories. Abundance trajectories and velocity vectors were transformed back to protein space from the PC-space before analysis.

Protein-wise velocity summaries were computed from the generated validation trajectories. The absolute velocity |*v*_*θ,p*_(*t, x*_*t*_)| was used to rank dynamic proteins, while signed velocities were used to examine the direction and timing of protein-specific changes. In the sepsis analysis, mean absolute velocity was summarized over tissue, sepsis model, protein and disease time. Rank stability was assessed across the five independently trained models by ranking proteins within each tissue–model combination and comparing rank distributions across seeds. Generated and observed protein abundances were compared after averaging within disease-time windows. Velocity-profile similarities between generated trajectories were computed from protein-by-time velocity profiles and visualized with UMAP. Model-specific velocity contrasts were calculated by subtracting the mean protein-wise velocity profile from each model-stratified profile. The time of peak velocity was defined for each protein and generated trajectory as the time point with the largest absolute velocity.

In the COVID-19 analysis, validation trajectories were clustered according to the similarity of their full protein-by-severity velocity profiles using agglomerative clustering on the pairwise Euclidean distance matrix. Cluster-specific analyses summarized mean absolute velocity over severity, ranked proteins by mean absolute velocity within each cluster, and visualized signed velocity profiles for the union of the top dynamic proteins. For visualization of observed and generated proteomic states, UMAP embeddings were fitted on observed samples and generated trajectory states were projected into the same embedding space.

### 8.6 Baseline comparison analyses

To compare flow-matching velocity with conventional summaries of protein change, we performed three baseline analyses on the sepsis data. First, samples were split by normalized disease time into early (*t <* 0.5) and late (*t ≥* 0.5) groups. For each protein, mean abundance was compared between the two groups using Welch’s *t*-test, and *p*-values were adjusted with the Benjamini– Hochberg procedure. The resulting log fold changes were compared with mean signed flow-matching velocities averaged over generated validation trajectories.

Second, empirical finite-difference velocities were computed from observed data. For each protein, tissue and sepsis model, mean abundance was calculated at each observed normalized time point. Adjacent time points *t*_*i*_ and *t*_*i*+1_ were then used to define

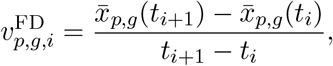

where *p* denotes protein and *g* denotes the tissue–model combination. Flow-matching velocities were averaged over generated trajectory points falling in the corresponding interval [*t*_*i*_, *t*_*i*+1_), yielding interval-matched estimates for comparison with 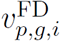 . Velocity roughness was quantified as the total absolute change in interval velocity divided by the spacing between interval midpoints,

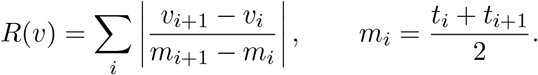

Third, a linear baseline was fitted separately for each protein, tissue and sepsis model using ordinary least squares,

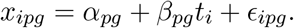

The fitted slope 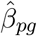 was treated as a constant linear velocity over the full disease trajectory. This provided a maximally smoothed reference against which both finite-difference velocities and flow-matching velocities were compared.

## 9 Author contributions

EH designed the project and conducted the analysis. EH wrote the first draft of the manuscript. JM supervised the project. Both EH and JM contributed to finalizing the manuscript.

## 10 Acknowledgements

We thank Jonas Wallin for insightful discussions.

## 11 Data availability

The COVID-19 datasets are available on the ProteomeXchange Consortium via the PRIDE partner repository with the identifier PXD025752 [33]. The source data that define the trajectories generated and analyzed in this study are available as tables in the accompanying GitHub repository. The raw data for the generated and analyzed trajectories are available on Zenodo with the identifier 10.5281/zenodo.20612314.

## 12 Code availability

The code is available on GitHub at https://github.com/ErikHartman/ProteoCFM. Notebooks in the repository contain the code necessary to generate trajectories as well as to perform downstream analysis and generate plots.

## 13 Supplementary information

### Differential abundance and regression coefficients as velocity summaries

Consider a single protein with abundance *x*_*p*_(*t*) along a temporal or pseudotemporal axis *t*. This quantity can be interpreted either as abundance along one modeled trajectory or, at the population level, as the conditional mean abundance *μ*_*p*_(*t*) = E[*X*_*p*_ | *t*]. Classical differential abundance compares two states, for example an initial state *t*_0_ and a later state *t*_1_. If the two states are separated by a fixed interval, then the same contrast can be written as a finite-difference rate of change,

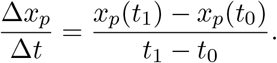

In many differential abundance analyses, Δ*t* is implicit because the comparison is simply “state 1 versus state 0”. Taking the log of *x*_*p*_ turns Δ*x*_*p*_ into the usual log fold change, and Δ*x*_*p*_*/*Δ*t* is the average log-abundance change per unit progression along the chosen axis.

In a linear regression analysis on the progression coordinate for observations *i* = 1, …, *n*, a protein-wise model

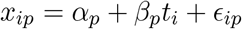

estimates the constant rate

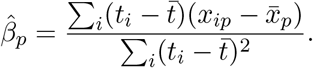

When observations occur at only two axis positions, *t*_0_ and *t*_1_, this slope reduces to the finite-difference estimate between the group means. More generally, 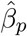 is the least-squares constant velocity for protein *p* over the observed range of the progression coordinate. On a log-abundance scale, it is a log fold-change rate per unit progression.

Proteome velocity is obtained by asking what happens when this rate is allowed to vary locally with time and proteomic state. Taking the limit as Δ*t →* 0 gives

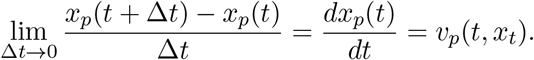

To estimate differential abundance from the velocity field, the velocity can be averaged:

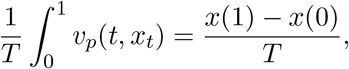

This relationship is shown in **Fig. 4a-b**.

Thus, differential abundance is a finite-difference summary over a chosen interval, whereas protein velocity is the corresponding instantaneous rate of change. This is the sense in which proteome velocity is a continuous, state-dependent analogue of differential abundance or a regression coefficient: differential abundance and *β*_*p*_ summarize net or average change over a chosen interval, while velocity resolves how that change is accumulated over the modeled trajectory.

**Figure S1:**
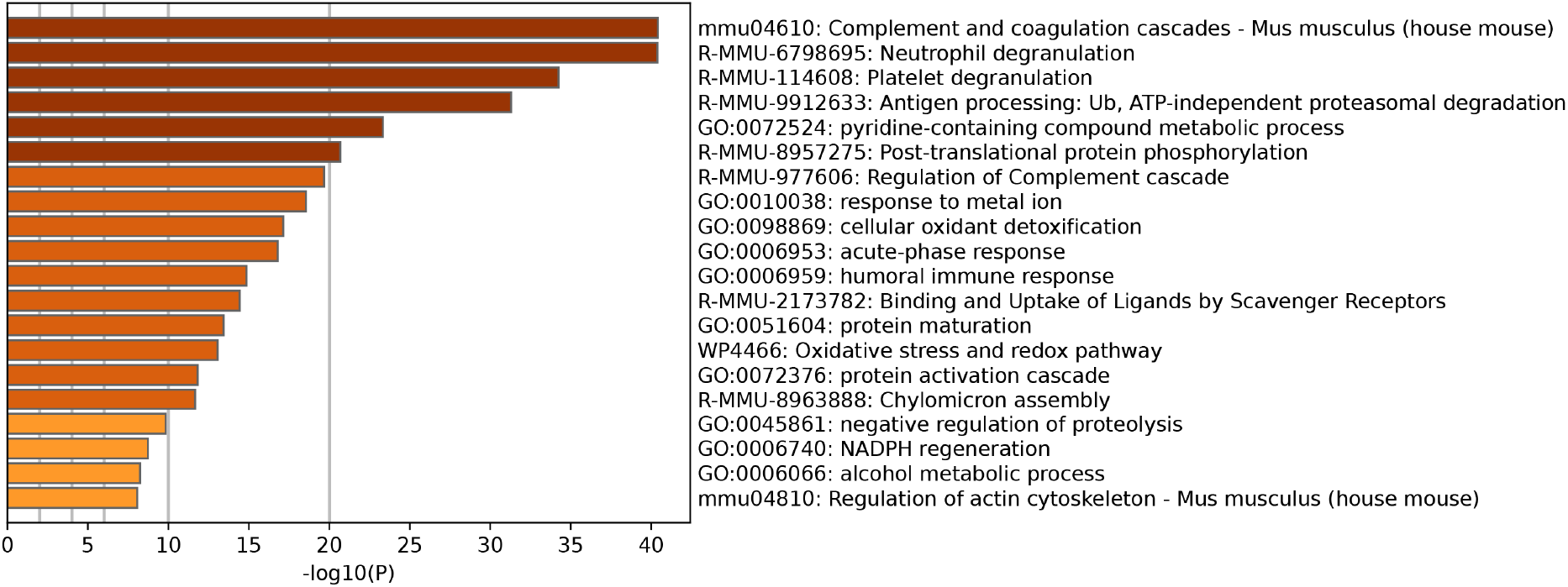
Biological pathway enrichment of the common sepsis proteins. Top 20 enriched terms across the sepsis proteome, colored by p-values. Created using Metascape [40].

**Figure S2:**
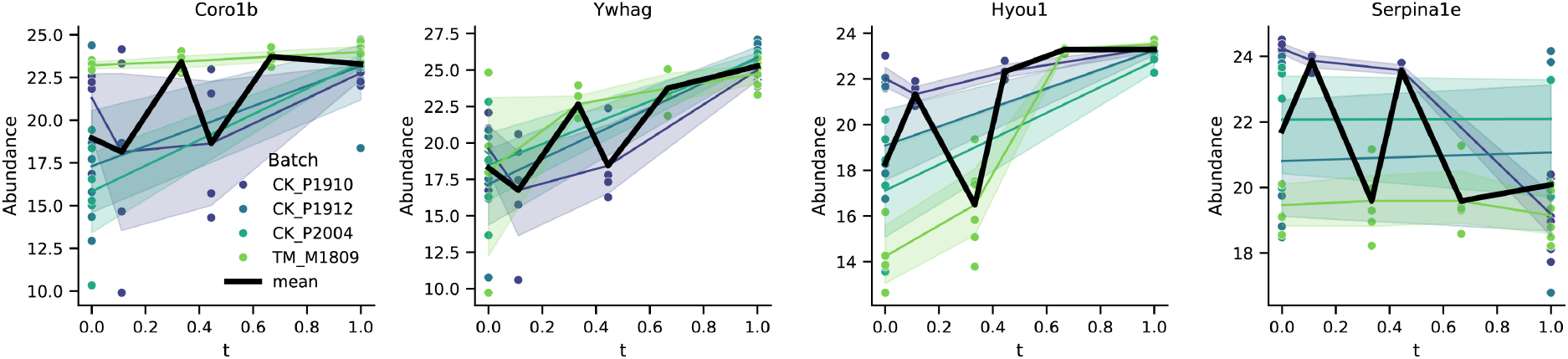
Abundance profiles for empirical velocity roughness outliers. Abundance over time of the proteins Coro1b, Ywhag, Hyou1 and Serpina1e which hade the highest finite difference velocity roughness.

